# CD8+CD20+ Cytotoxic T Lymphocytes Exhibit Augmented Degranulation and Pro-inflammatory Potential in Multiple Sclerosis

**DOI:** 10.64898/2026.01.29.702512

**Authors:** Özgür Albayrak, Selen Ünlü, Nazan Akkaya, Ali Burak Kızılırmak, Tansu Doran, Mina Üzülmez, Işıl Baytekin, Kemal Soylu, Mesrure Köseoğlu, Burcu Yüksel, Aysun Soysal, Atay Vural

## Abstract

The effectiveness of CD20-targeting therapies in multiple sclerosis (MS) underscores the role of B cells in the disease, yet the limited success of other B cell-specific treatments suggests additional mechanisms at play. Intriguingly, CD20 is also expressed on a subset of highly active memory T cells, particularly cytotoxic CD8+ T lymphocytes (CTLs). This study investigated the functional characteristics of CD8+CD20+ CTLs in MS. We observed a significant increase in CD8+CD20+ CTL prevalence in MS patients, with enhanced infiltration into the cerebrospinal fluid. Consistent with prior reports, these cells exhibited heightened proliferation and production of IFN-γ, TNF-α, and GM-CSF. Notably, we demonstrate for the first time that CD8+CD20+ CTLs display escalated degranulation and produce significantly higher levels of Granzyme B, Perforin, and Granzyme K compared to their CD20-counterparts, with further augmentation in pwMS compared to healthy controls. These findings suggest that in MS, CD8+CD20+ CTLs are actively recruited to the CNS, exhibiting enhanced cytotoxicity and a potent pro-inflammatory profile, particularly through elevated Granzyme K production, contributing significantly to the chronic inflammatory milieu and immunopathogenesis of MS.

## INTRODUCTION

CD20 is primarily expressed on B cells and targeting B cells with CD20 monoclonal antibodies has been a highly effective treatment for multiple sclerosis (MS). This effectiveness indicates that B cells play a crucial role in MS pathogenesis (1). However, it is intriguing that other B cell-specific therapies, such as atacicept and belimumab, have not shown the same effectiveness in treating MS (2). This disparity suggests that CD20 may influence MS pathogenesis through mechanisms beyond its expression on B cells.

In addition to B cells, CD20 is also found at low levels on a subset of T cells (3,4). Studies have shown that CD3+CD20+ lymphocytes exhibit an active phenotype and produce more cytokines when stimulated (4,5). These cells are primarily memory T cells and are enriched in type 1 cytotoxic CD8+ T cells (5). Furthermore, CD20+ T cells are also targeted by CD20 monoclonal antibody therapies, and their depletion has been shown to improve experimental autoimmune encephalomyelitis independently of B cells (6–8). Research indicates that CD3+CD20+ cells are significant in various autoimmune disorders, such as MS, rheumatoid arthritis, Sjögren’s syndrome, and psoriasis, and they play a role in cancer immunosurveillance (9,10).

Inflammatory processes activate CD8+ CTLs, leading to the secretion of inflammatory cytokines and the release of cytolytic granzymes and perforin through degranulation. While granzyme B is associated with cell death, granzyme K appears to be involved in blood-to-tissue migration and chronic inflammation, rather than cell death. A study in 2022 revealed a high presence of granzyme K expressing CD8+ CTLs in the synovial fluid of rheumatoid arthritis patients, with granzyme K enhancing the synthesis of proinflammatory cytokines and chemokines (11). Additionally, granzyme K induces IL-6 and IL-8 release from endothelial cells by activating the PAR-2 receptor (12). In post-mortem study of brains of pwMS, it was demonstrated that CD8+CD20+ CTLs were found abundantly in CD45RA-CD69+ memory cell compartment and exhibited effector properties, with granzyme K+ cells being particularly more abundant within this population compared to PBMCs, indicating their potential role in MS pathophysiology (13).

In this study, we primarily focused on CD8+CD20+ Cytotoxic T Lymphocytes (CTLs from now on) and their functional characteristics in MS and their contribution to MS immunopathogenesis. Our findings indicate that CD8+CD20+ CTLs are significantly more prevalent in MS and infiltrate cerebrospinal fluid (CSF) to a greater extent. It is known that CD8+CD20+ CTLs proliferate more and produce elevated levels of IFN-γ, TNF-α, and GM-CSF and our results are correlated with the existing literature. For the first time however, we identified that CD8+CD20+ CTLs show escalated degranulation and produce higher levels of Granzyme B, Perforin, and Granzyme K compared to CD8+CD20-cells, and that these characteristics are increased in pwMS compared to healthy controls. We showed that CD20+ CTLs are characterized by upregulated endogenous MS4A1 expression in pwMS relative to controls. Consistent with this distinct transcriptional profile, analysis of scRNA-seq datasets identified the MS4A1+ cytotoxic T cell population as belonging to the GZMK+ phenotype. These results indicate that in MS, a disease characterized by chronic inflammation, CD8+CD20+ CTLs migrate intensely into the cerebrospinal fluid with increased production of Granzyme K and demonstrate an active role in the immunopathogenesis of MS through enhanced cytotoxicity and proinflammatory cytokine production.

## MATERIALS and METHODS

### Study Population

This study was approved by the Koç University Clinical Research Ethics Committee under protocol number 2020.456.IRB1.169. A voluntary informed consent form was signed by all patients and healthy controls. 31 pwMS, 27 patients with other neurological disorders (OND) and 14 healthy controls (HC) were included in the study for comparison of CD8+CD20+ CTLs and 8 pwMS and 8 HCs were analyzed for functional studies. All patients received their diagnosis at Bakırköy Mazhar Osman Neuropsychiatry Hospital. 10ml of peripheral blood were drawn into vacutainer tubes containing K3EDTA. Cerebrospinal fluid (CSF) samples were obtained during a diagnostic lumbar puncturing. Samples were transported to Koç University Research Center for Translational Medicine for analysis. Clinical data including EDSS scores were obtained from patient charts. Patient and healthy control population included in the study is given in **Table 1**.

**Table 1.** Table shows the demographic data of subjects included in the Neuroinflammation Cohort. RRMS=Relapsing Remitting MS, GBS=Guillian-Barre Syndrome, MMN=Multifocal Motor Neuropathy, PCD=Paraneoplastic Cerebellar Degeneration, CIDP=Chronic Inflammatory Demyelinating Neuropathy, VSCT=Vasculitis, NBT=Neuro Behçet’s, HE=Herpes Encephelitis, ILAV=Inflammatory Lesions After Vaccination, IIH=Idiopatic Intracranial Hypertension, HA=Headache, CH=Cervical Herniation, ALS=Amytrophic Lateral Sclerosis

| Group (total n) | Diagnostic subgroups (n) | Age, years, (mean±SEM) | Sex, F:M |
| --- | --- | --- | --- |
| **MS (31)** | RRMS (31) | 33 ± 1.7 | 20:11 |
| **OND (27)** | GBS (3), CIDP (3), MMN (1), PCD (1), VSCT (1), NBT (1), HE (1), ILAV (1), IIH (6), HA (4), Migraine (3), CH (1), ALS (1) | 47.3 ± 2.8 | 21:6 |
| **HC (14)** | | 30.3 ± 1.3 | 9:5 |

### PBMC Isolation and Preparation of CSF Samples

PBMCs from peripheral blood were isolated using density gradient centrifugation method. Blood samples were diluted 1:1 with facs buffer (PBS containing 1% BSA), carefully layered on equal volume of Lymphoprep d=1.077g/ml (Axis-Shield, Norway) and centrifuged at 500g for 30 minutes at room temperature without brakes. Buffy coat containing PBMCs were collected into a sterile 50ml falcon tube and washed with equal volume of staining buffer at 500g for 5 minutes. After centrifugation, 1x10^7^ cells were cryopreserved in liquid nitrogen for subsequent functional analysis whereas 6x10^6^ cells were used freshly in immunophenotyping experiments. CSF samples were centrifuged at 400g for 15 minutes at 40C. After centrifugation, the cell pellet was used freshly for immunophenotyping studies.

### Immunophenotyping of PBMC and CSF samples

PBMC and CSF samples were incubated in the dark for 10 minutes with 100 µL of PBS containing Zombie NIR fixable viability dye (Biolegend, USA). Tubes were then centrifuged at 500g for 5 minutes with 2 mL of PBS. After centrifugation, all liquid was aspirated from the tubes, and 100 µL of staining buffer containing mouse anti-human CD3 FITC (Clone HIT3a), CD8 Alexa Fluor 700 (Clone SK1), CD20 Brilliant Violet 421 (except for the CD20 FMO tube, Clone 2H7), CD19 APC-Cy7 (Clone HIB19), and CD14 APC-Cy7 (Clone HCD16), antibodies were added to the samples. 16 PBMCs (pwMS n=8, Healthy Control n=8) were also stained with mouse anti human KLRG-1 PE (Clone SA231A2) and CD127 BV711 (Clone A019D5). All antibodies were acquired from Biolegend (Biolegend, USA). Samples were incubated in the dark at room temperature for 20 minutes. After incubation, tubes were centrifuged with 2 mL of staining buffer at 500g for 5 minutes and resuspended in 500 µL of staining buffer before being analyzed on Attune NxT flow cytometer (Thermo Scientific, USA). Flow cytometric analyses were performed using FlowJo version 10.10.0 (BD Biosciences, USA). CD8+CD20+ CTLs were identified using the gating strategy shown in **Figure S1**.

### Soluble Factors of CD8+CD20+ CTLs

Eight cryopreserved PBMCs from healthy controls were thawed at 37^°^C water bath and taken into complete medium (RPMI 1640 + 10% FBS + 1% Penicillin-Streptomycin, GIBCO, USA). All the centrifugation steps were performed at 500g for 5 minutes unless reported otherwise. PBMCs were transferred into 12x75 mm flow cytometry tubes, incubated with 100 µL of PBS containing Zombie NIR fixable viability dye (Biolegend, USA) for 10 minutes in the dark at RT and washed with 2 mL of PBS. Subsequently, 100 µL of staining buffer containing mouse anti-human CD3 FITC (Clone HIT3A), CD4 PE-Dazzle 594 (Clone OKT4), CD8 Alexa Fluor 700 (Clone SK1), CD20 Brilliant Violet 421 (Clone 2H7), CD19 APC-Cy7 (Clone HIB19), and CD14 APC-Cy7 (Clone HCD14) was added to the samples, followed by incubation for 20 minutes in the dark at room temperature. All antibodies were acquired from Biolegend (Biolegend, USA). After staining, the samples were washed with staining buffer, and CD8+CD20+ and CD8+CD20-cells were sorted using a FACS Aria III cell sorter. A 100 µm nozzle and a flow rate of ≤2000 events per second were used to ensure maximum purity and viability. After sorting, a portion of the sorted cells (∼1000 cells) were re-run to check the sorting purity and samples with >95 purity was used for experiments (**Figure S2)** After sorting CD20+ and CD20-CTLs were seeded into 96 well round bottom plates in 100 ul RPMI medium with or without Cell Activation Cocktail (PMA/Ionomycin without Brefeldin A, Biolegend, USA) and stimulated for 4 hours at 37^°^C 5% CO_2_ incubator. Following stimulation, plates were centrifuged, and supernatants were collected. Concentrations of Granzyme B, Granzyme A, Perforin, IFN-γ, TNF-α, sFAS, and sFASL (expressed in pg/mL) were measured using Legendplex Human CD8/NK Kit (Biolegend, USA) according to the manufacturer’s protocols. Measurements were performed on Attune NxT flow cytometer (Thermo Scientific, USA) and analysis were done using LegendPlex Data Analysis Software Suite (Biolegend, USA).

### Functional Analysis of CD8+CD20+ CTLs Between pwMS and Healthy Controls

Sixteen cryopreserved PBMCs (pwMS n=8, Healthy Control n=8) were thawed at 37^°^C water bath and taken into complete medium (RPMI 1640 + 10% FBS + 1% Penicillin-Streptomycin, GIBCO, USA). All the centrifugation steps were performed at 500g for 5 minutes unless reported otherwise. PBMCs were seeded into 96 well round bottom plates at a density of 1 x 10^6^ cells/100 ul RPMI medium with or without Cell Activation Cocktail (PMA/Ionomycin without Brefeldin A, Biolegend, USA) containing mouse anti-human CD107a Brilliant Violet 421 antibody (Clone H4A3, Biolegend, USA) and stimulated for 1 hour at 37^°^C 5% CO_2_ incubator. After 1 hour, Brefeldin A (Biolegend, USA) was added into the wells and additional stimulation was carried out for 3 hours. After stimulation, plates were centrifuged, supernatants were discarded, and cells were resuspended and incubated with PBS containing Zombie NIR fixable viability dye (Biolegend, USA) for 10 minutes at RT in dark. After incubation, plates were centrifuged, supernatants were discarded, and cells were resuspended with 100ul staining buffer containing CD3 FITC (Clone HIT3a), CD8 Alexa Fluor 700 (Clone SK1), CD20 APC (Clone 2H7), CD19 APC-Cy7 (Clone HIB19), and CD14 APC-Cy7 (Clone HCD14) antibodies and incubated at RT in dark for 20 minutes. After incubation, plates were centrifuged, and cells were fixed with Fixation Buffer and permeabilized by Permeabilization Wash Buffer (Biolegend, USA) according to the manufacturer’s instructions. After fixation/permeabilization, cells were resuspended with permeabilization buffer containing mouse anti-human granzyme B PE-Cyanine5 (Clone QA16A02), perforin PE (Clone B-D48), granzyme K PE-Cyanine7 (Clone GM26E7) for Panel 1, and IFN-γ PE (Clone B27), TNF-α PE-Cyanine7 (Clone MAB11), rat anti-human GM-CSF PerCP-Cyanine 5.5 (Clone BVD2-21C11) for Panel 2 and incubated for 20 minutes at RT in dark. After incubation, plates were washed with permeabilization buffer and staining buffer respectively, resuspended with staining buffer, transferred into 15x75mm tubes, and analyzed with Beckman Coulter Cytoflex SRT flow cytometer (Beckman Coulter, USA). Flow cytometric analysis was done with FlowJo v 10.10.0 (BD Biosciences, USA)

### MS4A1 Expression of CD8+CD20+ Cells

Twenty cryopreserved PBMCs (pwMS, n=10; Healthy Controls, n=10) controls were thawed at 37^°^C water bath and taken into complete medium (RPMI 1640 + 10% FBS + 1% Penicillin-Streptomycin, GIBCO, USA). All the centrifugation steps were performed at 500g for 5 minutes unless reported otherwise. PBMCs were transferred into 12x75 mm flow cytometry tubes, incubated with 100 µL of PBS containing Zombie NIR fixable viability dye (Biolegend, USA) for 10 minutes in the dark at RT and washed with 2 mL of PBS. Subsequently, 100 µL of staining buffer containing mouse anti-human CD3 FITC (Clone HIT3A), CD4 PE-Dazzle 594 (Clone OKT4), CD8 Alexa Fluor 700 (Clone SK1), CD20 PE (Clone 2H7), CD19 APC-Cy7 (Clone HIB19), and CD14 APC-Cy7 (Clone HCD14) was added to the samples, followed by incubation for 20 minutes in the dark at room temperature. All antibodies were acquired from Biolegend (Biolegend, USA). After staining, the samples were washed with staining buffer, and CD8+CD20+ and CD8+CD20-cells were sorted using a Cytoflex SRT cell sorter (Beckman Coulter, USA). A 100 µm nozzle and a flow rate of ≤2000 events per second were used to ensure maximum purity and viability. After sorting, a portion of the sorted cells (∼1000 cells) were re-run to check the sorting purity and samples with >95 purity was used for experiments. Total RNA was isolated from the sorted cells using the QIAGEN RNeasy Mini kit with minor modifications to the protocol. Briefly, the isolated cells were lysed using 1ml Trizol (Thermo-Scientific, USA), and 200 µl chloroform (Sigma-Aldrich, Germany) was added, followed by centrifugation at 12,000 x g for 10 minutes at 4°C. After centrifugation, the aqueous phase was transferred to a new tube, and an equal volume of 70% ethanol was added before proceeding with the kit protocol. Following the quantification of isolated RNA quality using a NanoDrop 2000, RNA was reverse transcribed into cDNA using the iScript cDNA synthesis kit (BioRad, USA). Subsequently, qPCR was performed using a QuantiStudio 7-Plex system with TaqMan B2M housekeeping gene and MS4A1 primer-probe sets (Thermo-Scientific, USA), both containing a 5’FAM reporter and a 3’MGB quencher. qPCR reactions were carried out with TaqMan Fast Advanced Master Mix (Thermo-Scientific, USA). Samples were run in duplicate, and comparative CT method (2 ^−ddCT^) was used for fold-change calculation between healthy controls and pwMS.

## Statistical Analysis

Statistical analyses were conducted using GraphPad Prism version 10.4.0. Kruskal-Wallis test was utilized to compare the percentages of CD8+CD20+ CTLs among pwMS, OND patients, and healthy controls and Mann-Whitney U test was employed to compare the population of these cells in the CSF between pwMS and OND patients. Paired Wilcoxon test was used to compare the frequencies of CD8+CD20+ CTLs between PBMC and CSF. Two-way ANOVA test with Bonferroni correction was performed to compare the functions of CD8+CD20+ and CD8+CD8+CD20-CTLs within the PBMCs of pwMS and healthy controls. Paired Wilcoxon test was used to compare the concentrations of cytokines and cytolytic molecules in CD20+ and CD20-CTLs. Descriptive statistics for the comparison of results were given as mean±SEM.

### scRNA-seq analysis

Peripheral blood single cell RNA-sequencing data from healthy twins, twins with subclinical neuroinflammation (SCNI) or twins with MS were obtained from Gene Expression Omnibus with GSE276167 accession number (14) and processed in Scanpy (15). The data contained negatively isolated CD8 T cells only. Data preprocessing and clustering performed by the original study were retained and MS4A1^+^ CD8 T cells were defined as those with nonzero MS4A1 expression (counts > 0). Expression of classical B-cell markers (BANK1, CD79A, CD79B, JCHAIN, CD19, CD40) and T-cell markers (CD3D/E/G, CD8A/B) were inspected to confirm true T cell identity of MS4A1^+^ subsets. Differential expression between MS4A1^+^ and MS4A1^−^ CD8 T cells was performed using Scanpy’s “sc.tl.rank_genes_groups” function, with Wilcoxon rank-sum statistics and Benjamini–Hochberg correction. Over-representation analysis (ORA) with Enrichr was performed via gseapy package in Python on the significantly differentially expressed genes from the CD8 MS4A1 comparison (upregulated and downregulated genes were analyzed separately, FDR < 0.05) (16,17). The following libraries were queried: GO Biological Process (2025), GO Cellular Component (2025), GO Molecular Function (2025), Reactome (2024), KEGG (2021 Human), MSigDB Hallmark (2020), BioCarta (2016), HDSigDB Human (2021), and GeneSigDB. Enrichment P values (Enrichr/Fisher’s exact) were Benjamini–Hochberg adjusted per library.

## RESULTS

### CD8+CD20+ CTLs are more abundant in pwMS, with higher levels observed in CSF samples

In the initial phase of our study, we investigated the impact of CD8+CD20+ CTLs on the immunopathogenesis of MS in the context of clinical parameters. Demographic and clinical characteristics of the study population is given in Table 1. The comparison of CD8+CD20+ CTLs within peripheral blood among pwMS, OND patients, and HCs revealed a significantly higher percentage of these cells in pwMS compared to other groups (pwMS=10.75±0.8395, OND=5.035±0.1291, HC=4.666±0.2548, p<0,0001). Similarly, examination of CSF samples from MS and OND patients, demonstrated an increased percentage of CD8+CD20+ CTLs in the CSF of MS patients (pwMS=15.55±0.7884, OND=7.481±0.4101). Furthermore, paired analysis of blood and CSF samples revealed that CD8+CD20+ CTLs were significantly more abundant in the CSF compared to the blood (Blood=7.919±0.5386, CSF=12.31±0.7872) **(Figure 1A and 1B)**.

**Figure 1.**
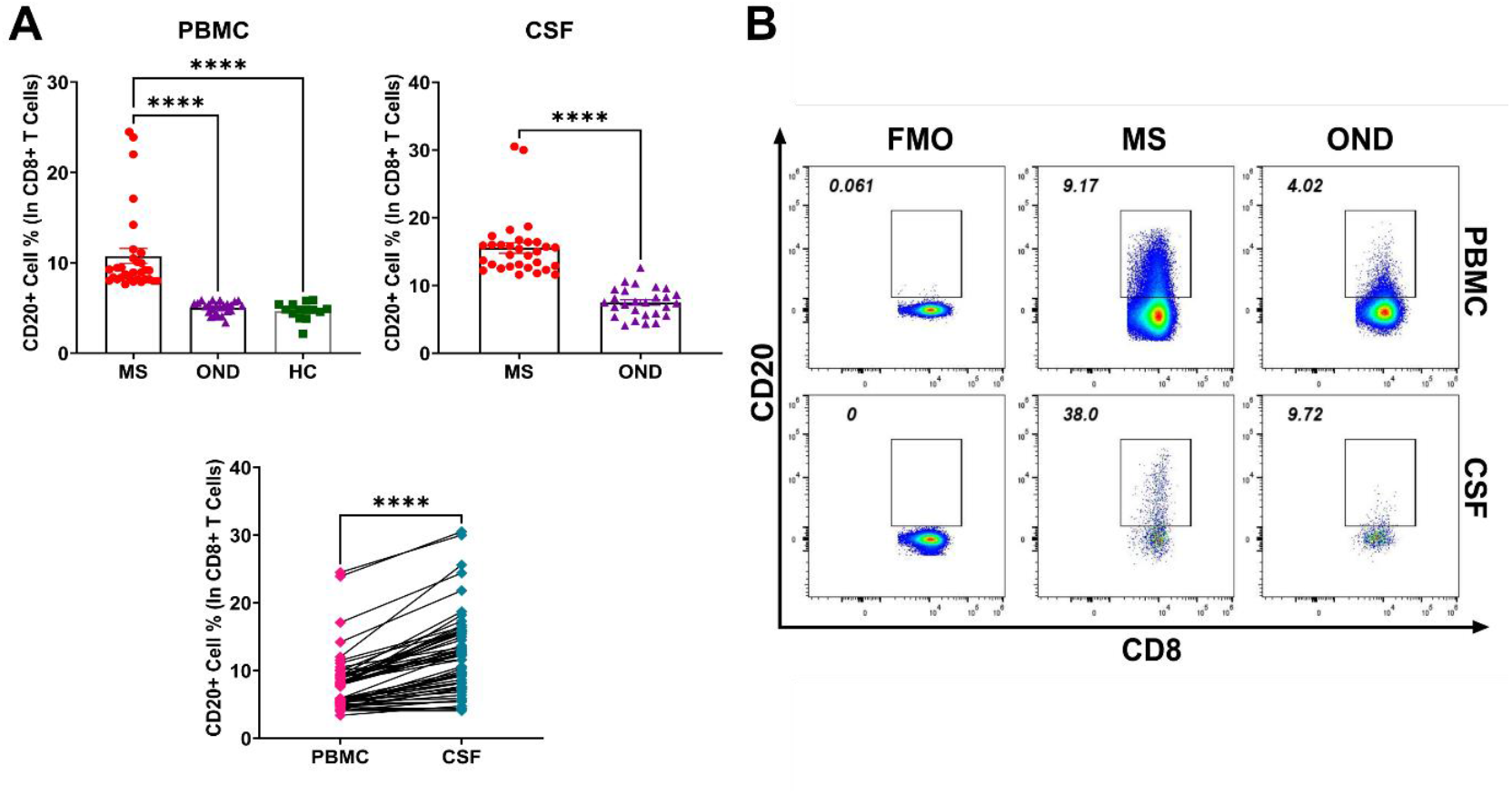
The figure illustrates comparisons of CD20+ CTLs in PBMCs and CSF samples. KruskallVallis Test with Dunn’s correction was used for multiple comparison analysis. Mann-Whitney U Test was used for independent samples and Wilcoxon Test was used for paired analysis. A) CD20+ CTLs were found to be significantly elevated in both PBMCs and CSF samples of MS patients. B) Representative flow cytometry graphic depicting the distribution of CD20+ CTLs between groups. ****p<0.0001, ***p<0.001, **p<0.01, *p<0.05.

Altogether, these results show that the CD8+CD20+ CTL population is increased in pwMS and ii) is enriched in the CSF compared to blood.

### CD8+CD20+ CTLs are More Cytotoxic, Produce Proinflammatory Cytokines and Cytolytic Molecules More Abundantly and Proliferate More Efficiently Than CD8+CD20-CTLs

In the second phase of our study, we aimed to determine i) compare the functional characteristics of CD20+ vs CD20-CTLs ii) whether CD8+CD20+ CTLs have a functional role in the immunopathogenesis of MS.

When we analyzed the soluble factors secreted by sorted CD20+ and CD20-cytotoxic T cells from healthy control PBMCs stimulated with PMA/I via Cytometric Bead Array, our analyses demonstrated that CD8+CD20+ CTLs released substantially greater amounts of Granzyme B, Granzyme A, Perforin, IFN-γ, TNF-α, IL-4, sFAS, and sFASL into the culture supernatant following stimulation (**Figure 2**, p<0.01).

**Figure 2.**
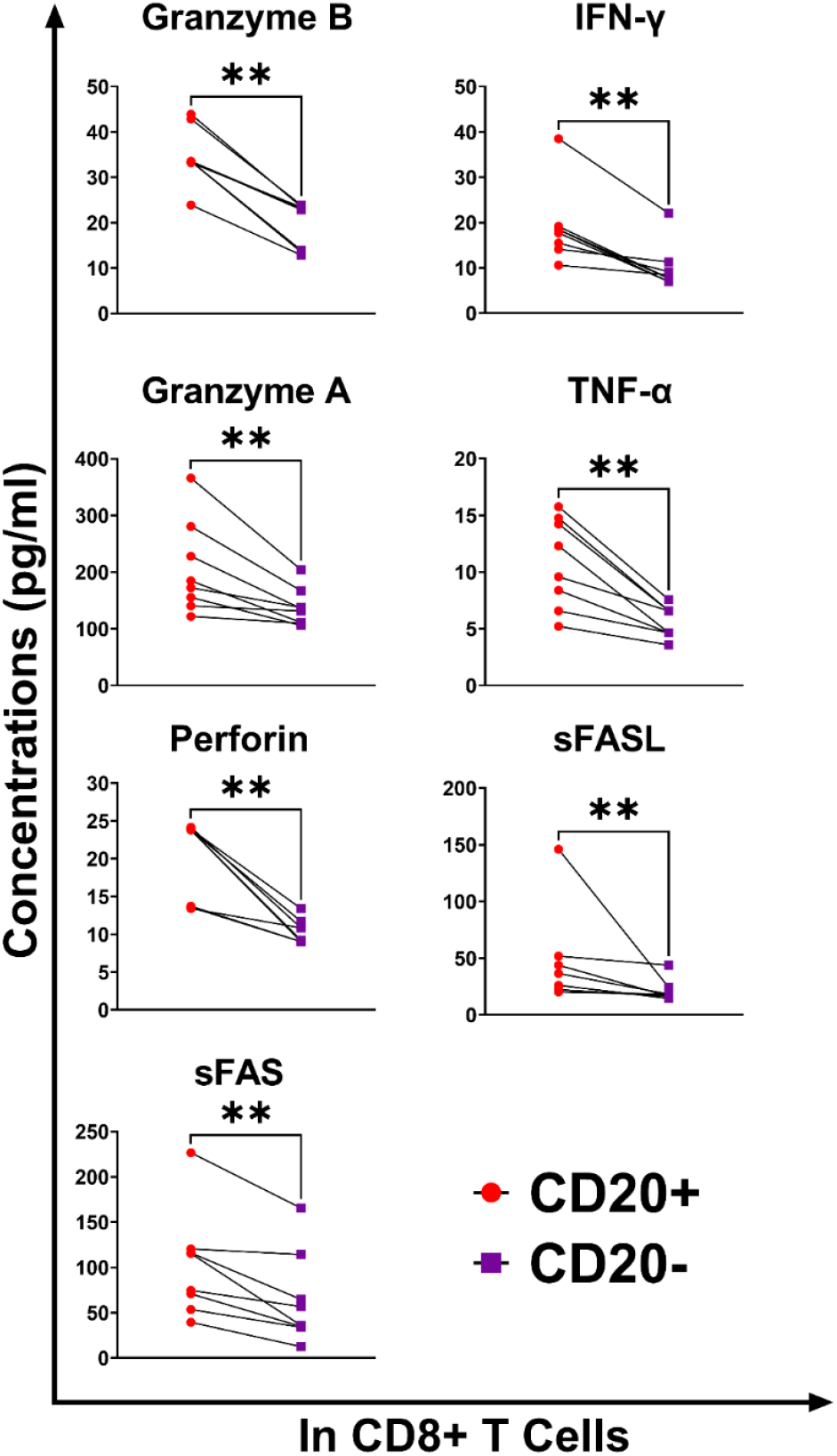
Figure shows the statistical analysis of soluble factors released by CD8+CD20+ and CD8+CD20-cells, after FACS sorting and PMA/I stimulation. Concentrations were measured via cytometric bead array and calculated as pg/ml. Paired Wilcoxon test was used for statistical analysis. ****p<0.0001, ***p<0.001, **p<0.01, *p<0.05

Subsequently, we compared the proliferation capacities of CD20+ and CD20-CTLs via T cell proliferation assay. When analyzing total proliferation, we demonstrated that CD20+ CTLs exhibited significantly elevated proliferation compared to CD20-cells (CD20+ CTLs 90.3 ± 1.3%, CD20-CTLs 71.8 ± 1.7, p<0.01, **Figure S2A and S2B**) In sub-analyses where CFSE peaks were evaluated as distinct cell generations, we observed that the non-dividing fraction and the less proliferative 1st and 2nd generations had a higher frequency of CD20-cells compared to CD20+ cells. However, in the more proliferative 3rd, 4th, and 5th generations, CD20+ CTLs were found to be substantially more frequent (p<0.01, **Figure S2C and S2D**)

Next, we analyzed total PBMCs of pwMS and healthy controls stimulated with PMA/I. Our analyses revealed that CD8+CD20+ CTLs exhibit a significantly abundant percentage of CD107a+ degranulated cells compared to CD8+CD8+CD20-cells, in both pwMS and healthy controls (pwMS p<0.0001, HC p<0.01). When comparing mean fluorescence intensity (MFI) values, we observed that CD8+CD20+ CTLs expressed higher levels of CD107a on their surface compared to CD8+CD8+CD20-cells, in both pwMS and healthy controls (pwMS p<0.0001, HC p<0.05). Upon comparing the CD8+CD20+ CTLs of pwMS healthy controls, we found that CD8+CD20+ CTLs from MS patients exhibited a higher degranulation potential, as indicated by both a greater percentage of CD107a+ cells (p<0.01) and higher CD107a MFI values (p<0.001) However, we did not identify a significant difference in degranulation potential when we compared CD8+CD20-CTLs between pwMS and healthy controls. (**Figure 3A and 3B**). Subsequently, to further investigate the cytotoxic potential of CD8+CD20+ CTLs, we compared Granzyme B+, Perforin+ cell frequencies and the expression levels of these molecules based on their MFI values within CD8+CD20+ and CD8+CD8+CD20-CTLs between pwMS and healthy controls (Representative flow cytometry gating figures in **Figure S2)**. In our analyses, when comparing CD8+CD20+ and CD8+CD20-CTLs in terms of Granzyme B and Perforin, we found no significant difference within healthy controls (p>0.05). In pwMS however, we observed significantly elevated frequencies of Granzyme B+ and Perforin+ cells in CD8+CD20+ CTLs (Granzyme B p<0.0001, Perforin p<0.001). When examining MFI values, we found that both Perforin and Granzyme B expressions were elevated in the CD8+CD20+ CTLs of both pwMS (Granzyme B p<0.001, Perforin p<0.01) and healthy controls (Granzyme B p<0.0001, Perforin p<0.0001). Subsequently, we aimed to determine the functional contribution of CD8+CD20+ CTLs to the immunopathogenesis of MS by comparing their cytotoxic potential between pwMS and healthy controls. At the conclusion of our analyses, we observed the following as Granzyme B and Perforin molecules were significantly escalated as in both percentage and MFI values within the CD8+CD20+ CTLs of pwMS compared to healthy controls (Granzyme B+% p<0.001, MFI p<0.01; Perforin+% p<0.001, MFI p<0.001) whereas no significant changes were observed in the CD8+CD20-compartment (p>0.05). In the literature, Granzyme K is shown to be associated with CD8+CD20+ CTLs and is described as a molecule primarily responsible for the transition from blood to tissue rather than cytotoxicity (12, 13). In our study, we investigated the Granzyme K molecule and observed a significant elevation in both cell frequency and MFI values within the CD8+CD20+ CTLs of both pwMS and healthy controls compared to CD8+CD20-cells (Cell % p<0.0001, MFI p<0.0001). Analysis within CD8+CD20+ CTLs of pwMS and healthy controls revealed abundant frequency of Granzyme K+ cells (p<0.001) and elevated Granzyme K MFI values (p<0.0001) in pwMS compared to healthy controls. However, we did not observe a significant difference between the CD8+CD20-CTLs of pwMS and healthy controls. Furthermore, in our analysis of degranulated CD8+CD20+ cells, we also identified a larger population of cells producing granzyme B, perforin, and granzyme K. This increase was evident both in comparison to CD20-cells and within the CD20+ cells of pwMS relative to those of healthy controls. (p>0.05, **Figure 3C**).

**Figure 3.**
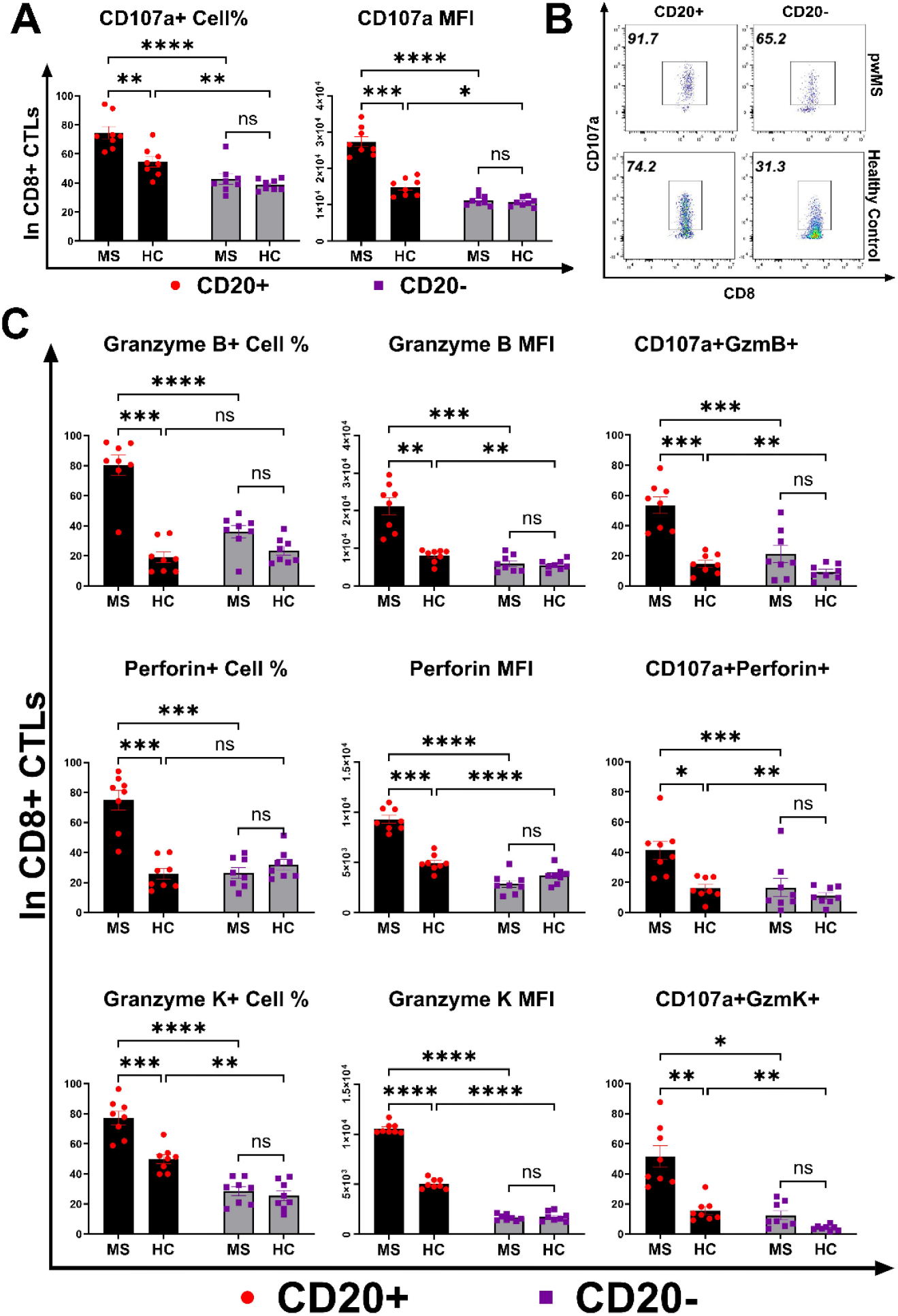
Figure represents the comparison of the cytotoxic potential of CD20+ and CD20-CTLs between MS patients and healthy controls. Two-way ANOVA with Bonferroni correction was used for statistical analysis. Frequencies of CD107a+ cells and CD107a expression were significantly elevated in CD20+ CTLs of both pwMS and healthy controls compared to CD20-CTLs. A) The cytotoxic potential of CD20+ CTLs of pwMS were found significantly greater than CD20+ CTLS of healthy controls. However, there were no significant difference between CD20-CTLs of pwMS and healthy controls regarding cytotoxic potential. B) Representative gating for CD20+ and CD20-CTLs of pwMS and Healthy Controls. C) Granzyme B+, Perforin+ and Granzyme K+ cell frequencies as well as MFI values of these molecules were found greatly escalated in both resting and degranulated (CD107a+) CD20+ CTLs of pwMS. ****p<0.0001, ***p<0.001, **p<0.01, *p<0.05.

To investigate the cellular immunity-based contribution of CD8+CD20+ CTLs to the MS immunopathogenesis, we compared the IFN-γ+, TNF-α+, and GM-CSF+ cell frequencies and MFI values of these cytokines between CD8+CD20+ and CD8+CD20-CTLs of pwMS and healthy controls. In the post-PMA/I stimulation analysis, we observed that the percentages of IFN-γ+, TNF-α+, and GM-CSF+ cells, as well as the MFI values of these cytokines, were significantly elevated within the CD8+CD20+ CTL population compared to CD8+CD20-CTLs in both pwMS (IFN-γ+ % p<0.001, MFI p<0.001; TNF-α+ % p<0.01, MFI p<0.001; GM-CSF+ % p<0.0001, MFI p<0.01) and healthy controls (IFN-γ+ % p<0.01, MFI p<0.01; TNF-α+ % p<0.01, MFI p<0.001; GM-CSF+ % p<0.01, MFI p<0.05).

Subsequently, we compared the percentages of cells producing aforementioned cytokines and their MFI values between the CD8+CD20+ CTL populations of pwMS and healthy controls. Our analyses indicate that the percentages of IFN-γ+, TNF-α+, and GM-CSF+ cells, as well as their expression levels based on MFI values, are significantly elevated in the CD8+CD20+ CTLs of pwMS compared to healthy controls (IFN-γ+ % p<0.01, MFI p<0.05; TNF-α+ % p<0.01, MFI p<0.001; GM-CSF+ % p<0.001, MFI p<0.001), whereas no significant differences were observed among CD8+CD20-CTLs (p>0.05). In the final phase, we compared the percentages of cells producing IFN-γ, TNF-α, and GM-CSF within the CD107a+ degranulated cell population between CD8+CD20+ and CD8+CD20-CTLs of MS patients and healthy controls. Our analyses revealed that i) CD107a+CD8+CD20+ CTLs have significantly higher cytotoxic potential and cytokine production compared to CD8+CD20-cells, and ii) the CD8+CD20+ CTLs of pwMS exhibit much greater cytotoxic potential and cytokine production compared to those of healthy controls. However, no significant differences were observed when analyzing the CD8+CD20-cells of pwMS and healthy controls (**Figure 4A, Figure S3**).

**Figure 4.**
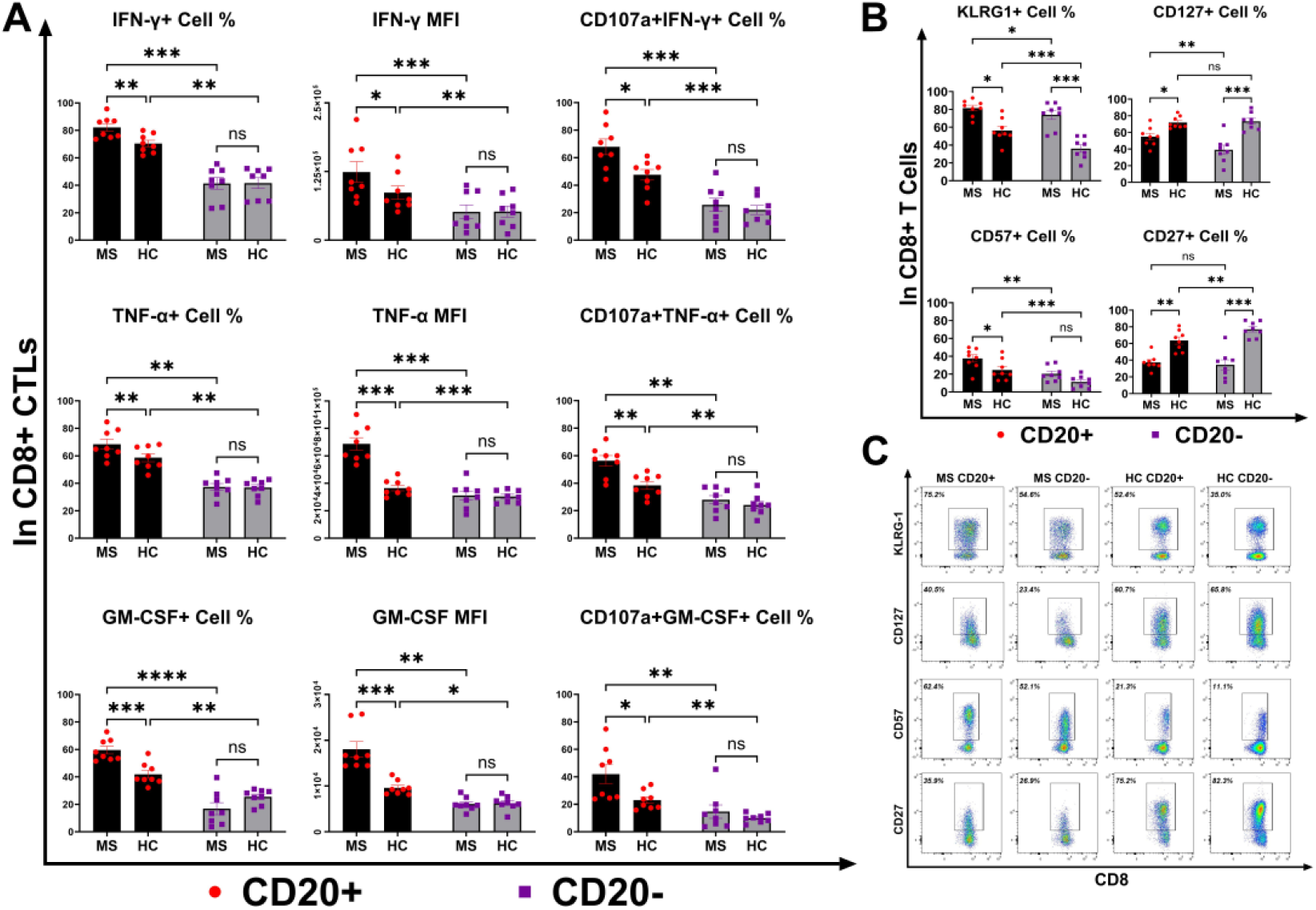
Figure shows the comparison of the Granzyme B+, Perforin+, Granzyme K+, IFN-γ+, TNFα+ and GM-CSF+ cell frequencies of degranulated (CD107a+) CD20+ and CD20-CTLs between MS patients and healthy controls. Two-way ANOVA with Bonferroni correction was used as statistical analysis method. Elevated cytotoxicity and cytokine production was observed in CD20+ CTLs of both pwMS and healthy controls compared to CD20-CTLs. Cytotoxic potential of CD20+ CTLs of pwMS were found significantly greater than CD20+ CTLs of healthy controls. However, there were no significant difference between CD20-CTLs of pwMS and healthy controls. ****p<0.0001, ***p<0.001, **p<0.01, *p<0.05.

### Upregulation of MS4A1 Gene Expression in CD20+ Cytotoxic T Cells of Patients with Multiple Sclerosis

To assess transcriptional changes of MS4A1 gene in CD20+ CTLs, we isolated these cells from PBMCs of healthy controls and pwMS via FACS. Post-sort analysis confirmed high purity of the isolated populations prior to RNA extraction (**Figure 5A**). Following cDNA synthesis, MS4A1 gene expression levels were quantified using the 2^^-ddCT^ method. As shown in **Figure 5B**, MS4A1 mRNA levels were significantly upregulated in pwMS compared to HCs (p < 0.0001). To validate the concordance between transcriptional and translational expression, we analyzed the correlation between MS4A1 dCT values (B2M - MS4A1) and surface CD20 protein expression. We observed a robust positive correlation between MS4A1 gene expression and both the frequency of CD20+ CTLs (r = 0.8074, p < 0.0001) and the MFI of CD20 (r = 0.8481, p < 0.0001) (**Figure 5C**).

**Figure 5.**
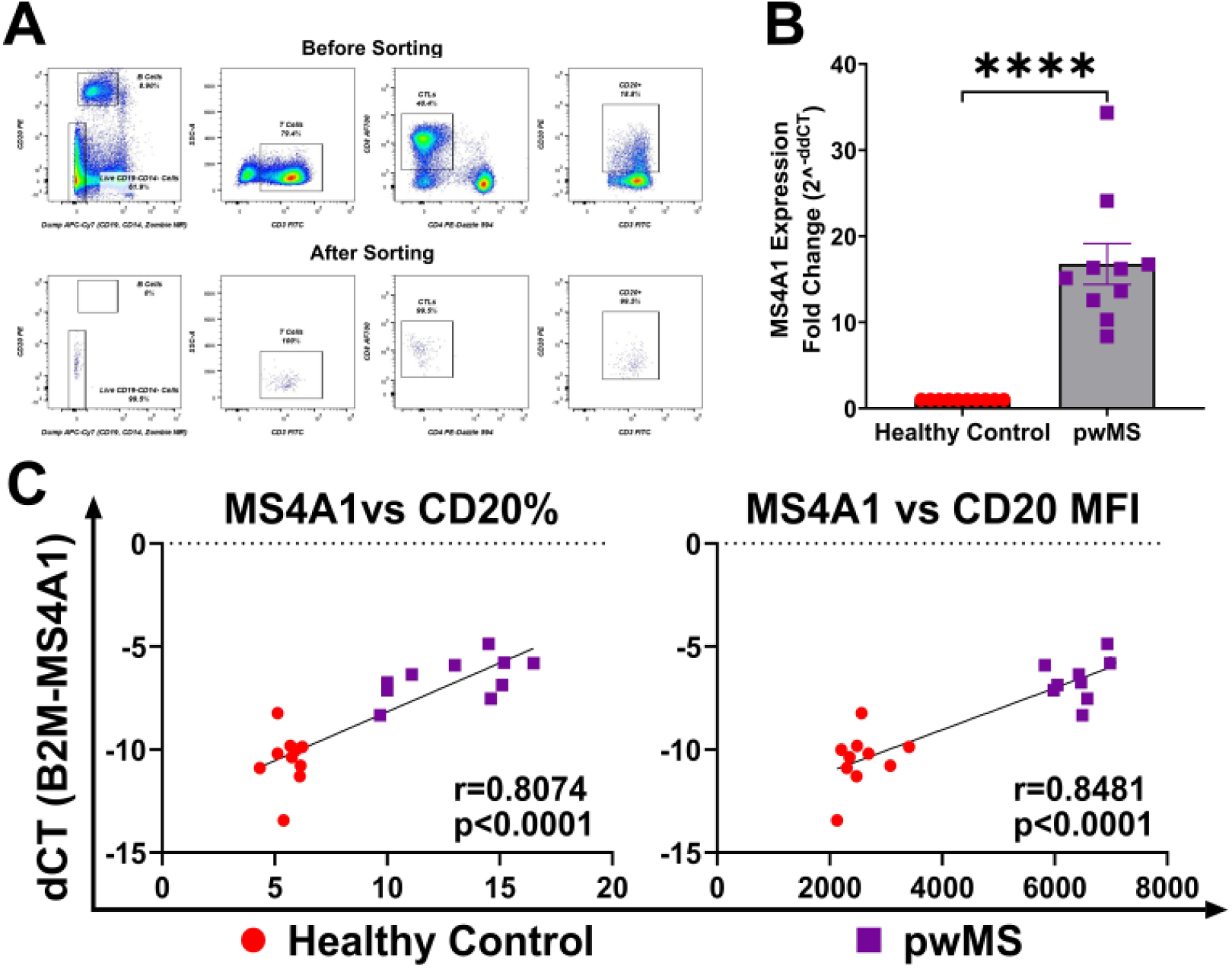
Enhanced MS4A1 gene expression in CD20+ cytotoxic T cells correlates with surface protein levels in pwMS. (A) Representative flow cytometry gating strategy used for the isolation of CD20+ cytotoxic T CTLs. (B) Relative MS4A1 mRNA expression in sorted CD20+ CTLs from healthy controls and pwMS. Gene expression was normalized to B2M, and fold change was calculated using the 2 ^-ddCT^ method. Results are given as mean ± SEM. Statistical significance was determined using the Mann-Whitney U test. (C) Pearson correlation analysis showing the relationship between MS4A1 gene expression (represented as dCT: B2M - MS4A1) and surface CD20 expression levels. The left panel correlates gene expression with the percentage of CD20+ cells (r=0.8074), and the right panel correlates gene expression with CD20 MFI (r=0.8481). ****p<0.0001, ***p<0.001, **p<0.01, *p<0.05.

**Figure 6.**
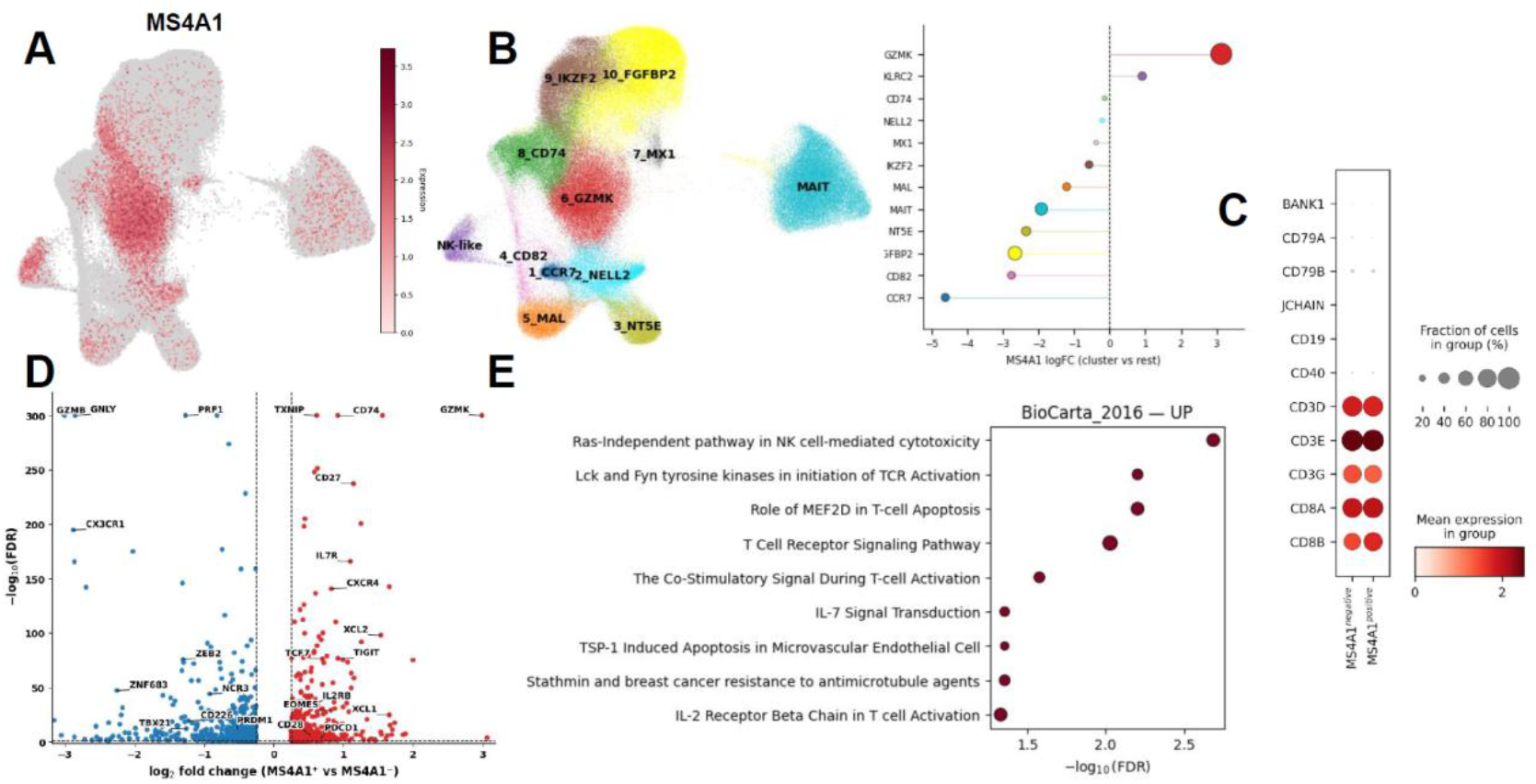
Single-cell transcriptomic profiling reveals a distinct GZMK+ activated phenotype in MS4A1+ CD8+ T cells. (A) UMAP feature plot visualizing the expression distribution of MS4A1 across the total CD8+ T cell landscape. Color intensity corresponds to the expression level. (B) Unsupervised clustering of CD8+ T cells visualized via UMAP (left). The lollipop plot (right) displays the log fold change (logFC) of MS4A1 expression across identified clusters, demonstrating significant enrichment specifically within the GZMK+ cluster (Cluster 6) compared to other subsets. (C) Dot plot validation of cell lineage markers. Expression of canonical B cell (CD19, CD79A, CD79B, BANK1) and T cell (CD3D, CD3E, CD8A, CD8B) genes is compared between MS4A1-negative and MS4A1-positive groups. The absence of B cell markers in the MS4A1+ group confirms that MS4A1 expression is intrinsic to T cells and not a result of B-cell doublet contamination. Dot size represents the fraction of expressing cells; color intensity indicates mean expression. (D) Volcano plot illustrating differentially expressed genes (DEGs) between MS4A1+ and MS4A1-CD8+ T cells. Red points denote significantly upregulated genes (e.g., GZMK, CD74, CXCR4, IL7R), while blue points indicate downregulated cytotoxic markers (e.g., GZMB, GNLY, PRF1). (E) Pathway enrichment analysis (BioCarta 2016) of genes upregulated in MS4A1+ cells. The analysis highlights a significant enrichment in T cell receptor (TCR) signaling and activation-related pathways.

### Single-Cell RNA Sequencing Identifies MS4A1+ CD8+ T Cells are Highly Enriched in the GZMK+ CD8+ T Cell Subset and Enriched for TCR Signaling Pathways

We evaluated MS4A1 expression across the 12 CD8 T cell clusters defined by the original study based on predominant gene expression patterns. MS4A1^+^ cells were most abundant in the GZMK cluster (17.44%), followed by the NK-like cluster (8.43%), while the naive-like CCR7 cluster contained the lowest proportion of MS4A1^+^ cells. To assess whether MS4A1^+^ CD8 T cell phenotypes differed across clinical groups (healthy, SCNI, MS), we visualized their distribution by UMAP and compared gene expression signatures via heatmap. MS4A1^+^ cells showed similar distribution across all three conditions, and their differentially expressed genes were largely concordant regardless of disease state. Given this consistency and to maximize statistical power, we performed a unified differential expression analysis comparing all MS4A1^+^ versus MS4A1^−^ CD8 T cells. This analysis revealed a gene expression signature consistent with GZMK^+^ CD8 T cell identity.

GZMK was markedly enriched in MS4A1^+^ CD8 T-cell fraction alongside upregulation of other granzymes GZMA and GZMM. Activation of granule machinery was further evident by the upregulation of LYST and RAB37, which mediate granule maturation and exocytosis (18,19). CST7 upregulation, which modulates granzyme activation, showed that granzyme activity is tightly regulated in those cells (20).

MS4A1^+^ CD8 T cells had upregulated chemokines (CCL5, XCL1, XCL2) and inflammatory cytokines (IL16, IL32) that can recruit and activate other immune cells at sites of inflammation (21–24). Among upregulated genes, there were a wide range of chemokine receptors and adhesion molecules that could allow diverse trafficking patterns: lymphoid homing via CCR7, SELL, and GPR183 for memory maintenance and antigen surveillance; lymphoid exit via S1PR1 (25); tissue infiltration via CXCR3, ITGA4, PECAM1 (26); peripheral tissue surveillance through CCR4 (27); and access to bone marrow via CXCR4 (28). Upregulation of CXCR3, CXCR4, CCL5 and ITGA4 were previously reported for GZMK+ CD8 T cells, in line with GZMK-biased identity of MS4A1+ CD8 T cells (29,30). MS4A1+ population has many upregulated MHC class II molecules (HLA-DPA1/DPB1, HLA-DQA1/DQB1, HLA-DRB1, HLA-DMA) and invariant chain CD74; which could indicate recent or sustained activation, as reported previously for GZMK+ CD8 T cells (11,14,31). The activated profile is further evident in upregulation of genes related to TCR signaling machinery. Those genes include Src kinases (LCK, FYN); adaptor proteins [LAT, LCP2 (SLP-76), SH2D1A (SAP), FYB1 (ADAP/SLP-130), TRAT1]; AP-1 family members (JUN, JUNB); PI3K/AKT pathway components and regulators (PIK3R1, AKT3, PIK3P1, INPP4B). Complementing this signaling machinery, many costimulatory molecules were prominent among upregulated genes including CD27, CD28, CD2, TNFSF8 and SLAM family members such as CD48, CD84, SLAMF6, SLAMF7, LY9. MS4A1^+^ CD8 T cells had upregulated death receptors FAS (CD95) and TNFRSF10A (TRAIL-R1), indicating susceptibility to activation-induced cell death (32). However, concurrent upregulation of anti-apoptotic proteins BCL2, BCL2A1, and BIRC3 suggests active survival signaling that may protect these cells from premature apoptosis (33). This balance between pro- and anti-apoptotic signals can maintain functional capacity of CD8 T cells (34). MS4A1^+^ CD8 T cells have upregulated IL7R and IL2RB, which could enable responsiveness to the IL-7 and IL-15 respectively. These cytokines are critical for memory T cell survival, maintenance, and homeostatic proliferation and are characteristic of long-lived memory populations (35). Additionally, upregulation of IL10RA indicates responsiveness to anti-inflammatory IL-10 signaling, which could prevent excessive activation (36). MS4A1^+^ CD8 T cells exhibit a distinctive transcription factor profile characterized by co-upregulation of TCF7 and EOMES. TCF7 maintains stem-like properties including self-renewal capacity, memory maintenance, and responsiveness to homeostatic cytokines (37–39) while EOMES drives cytotoxic and memory-like effector differentiation and was previously reported as a potential regulator of GZMK+ CD8 T cells (29). This TCF7+ EOMES+ phenotype was previously reported for GZMK+ CD8 T cells (40).

## DISCUSSION

In this study, we investigated the relationship between CD8+CD20+ CTLs and MS immunopathogenesis, as well as their functional properties. In our neurological diseases cohort, our findings indicated that i) the percentage of CD8+CD20+ CTLs was significantly increased in the PBMC and CSF samples of pwMS compared to those diagnosed with other neurological diseases and PBMCs of healthy controls, and ii) there was a positive correlation between the population of CD8+CD20+ CTLs and EDSS scores in both PBMC and CSF samples of pwMS. Our results are consistent with the studies conducted by Holley et al. and Von Essen et al., who also found elevated percentages of CD8+CD20+ CTLs in the PBMC and CSF samples of multiple pwMS (5,41). Additionally, Von Essen et al. identified a positive correlation between the CD8+CD20+ CTLs in PBMC and CSF samples EDSS scores in pwMS with RRMS (5). In another study, Von-Essen and colleagues demonstrated that intrathecal CD8+CD20+ CTLs were increased in the cerebrospinal fluid samples of PPMS patients and contributed to demyelination by showing a positive correlation with the presence of MBP (42).

Consistent with our findings, the significantly higher levels of granzyme K, which is known to be expressed highly in tissue migrating T cells (11,43), in CD8+CD20+ CTLs of MS patients compared to healthy controls, can explain why these cells are found in elevated numbers in the cerebrospinal fluid samples of MS patients.

In recent years, CD8+CD20+ CTLs have been extensively studied, particularly in the context of MS, as they are hypothesized to contribute to MS immunopathogenesis.

In a study by Ochs et al., it was shown that CD8+CD20+ CTLs possess pathogenic characteristics, and their elimination in mice resulted in a reduction of EAE (8). In another study by Quendt et al., it was demonstrated that CD8+CD20+ T cells exhibit a phenotype consistent with pathogenicity via pathogenic effector functions (44). Sabatino et al. also demonstrated the presence of myelin specific CD8+CD20+ CTLs in the peripheral blood of pwMS and showed that these cells are depleted with anti-CD20 therapy (45). In two studies conducted on post-mortem brains of pwMS, it was demonstrated that CD8+CD20+ CTLs are consistent with the CD103+CD69+ Trm phenotype and possess a dense population of Granzyme K+ and Granzyme B+ cells (13,46). However, our findings show that circulating CD8+CD20+ CTLs cells are both increased in proportion in both PBMCs and CSFs of pwMS and have become functionally more cytotoxic in pwMS than healthy controls. Therefore, the source of the perivascular CD8+CD20+CD69+ T cells in the brain may also be peripheral circulation.

A 2021 study using PBMC samples from PPMS patients demonstrated that the populations of cells producing Granzyme B and Perforin were elevated in CD8+CD20+ T cells (47). In our study, when comparing the CD8+CD20+ and CD8+CD20-CTLs of healthy controls, we found that the populations of Granzyme B+ and Perforin+ cells did not differ between CD8+CD20+ and CD8+CD20-cells; however, the MFI values were significantly higher in CD8+CD20+ cells. Conversely, in CD8+CD20+ CTLs compared to CD8+CD20-cells, we observed a marked increase in both the Granzyme K+ population and Granzyme K MFI values. When comparing pwMS to CD8+CD20-CTLs of pwMS and CD8+CD20+ CTLs of healthy controls, we identified a substantial increase in the populations of Granzyme B+, Perforin+, and Granzyme K+ CD8+CD20+ CTLs in CD8+CD20+ CTLs of pwMS.

In this study for the first time in the literature, we aimed to observe the cytotoxicity of CD8+CD20+ CTLs by testing CD107a degranulation. In the first phase of our study, we compared the CD107a degranulation of CD8+CD20+ and CD8+CD20-CTLs in PBMCs from pwMS and healthy controls. Our analyses revealed that a significantly higher percentage of CD8+CD20+ CTLs were CD107a+ compared to CD8+CD20-cells. Additionally, within the CD107a+ population, the percentages of Granzyme B+, Perforin+, and Granzyme K+ cells were significantly higher in CD8+CD20+ cells compared to CD8+CD20-cells. Interestingly, when comparing CD8+CD20+ and CD8+CD20-cells in healthy controls, we found no differences in the percentages of Granzyme B+ and Perforin+ cells, only higher MFI values in CD8+CD20+ cells, with a significant increase in both values observed only for Granzyme K. However, in pwMS, CD8+CD20+ CTLs showed substantial increases in both the percentages and MFI values of Granzyme B, Perforin, and Granzyme K. These results made us think that CD8+CD20+ CTLs may play a significant direct role in MS immunopathogenesis. To test this hypothesis, we compared the CD8+CD20+ CTLs of pwMS with those of healthy controls and found that the CD8+CD20+ CTLs in pwMS were significantly more degranulated and had a much higher population of Granzyme B+, Granzyme K+, and Perforin+ cells within the CD107a+ compartment compared to healthy controls. We have also found that CD8+CD20+ CTLs were produce IFN-γ, TNF-α, and GM-CSF more abundantly than CD8+CD20-CTLs, with significantly higher frequencies of these cytokine-producing cells present in the CD8+CD20+ population compared to both CD8+CD20-cells and the CD8+CD20+ CTLs of healthy controls. The lack of a significant difference in the cytotoxic potential and cytokine production of CD8+CD20-CTLs between pwMS and healthy controls further supports our hypothesis that the increased cytotoxic T cell functions in pwMS are most likely derived from CD8+CD20+ CTLs. Altogether our results suggest that a heightened functional capacity of CD8+CD20+ CTLs in terms of enhanced cytotoxicity and cytokine production, particularly in the context of MS, where this enhanced response may reflect their involvement in the disease’s immunopathogenesis.

The limitations of our study include the absence of cytotoxicity studies demonstrating that CD8+CD20+ CTLs kill target cells more efficiently than CD8+CD20-CTLs and the lack of molecular mechanism studies. However, despite these limitations, our results suggest with the utmost probability and possibility that CD8+CD20+ CTLs i) infiltrate the cerebrospinal fluid more effectively due to the high Granzyme K+ population, ii) contribute to MS immunopathogenesis by causing cell death through increased Granzyme B and Perforin mediated cytotoxicity with elevated CD107a degranulation and iii) escalate the inflammation through abundant IFN-γ, TNF-α, and GM-CSF production, all of which leading to demyelination. We suggest that more comprehensive studies should be conducted to examine the direct impact of CD8+CD20+ CTLs on demyelination and to investigate their contribution to MS immunopathogenesis.

## List of Supplementary Materials

Figure S1 to S3

## Acknowledgements

We would like to thank the patients and healthy volunteers who participated in this research project. The authors gratefully acknowledge the use of the services and facilities of the Koç University Research Center for Translational Medicine (KUTTAM), funded by the Presidency of Turkey, Presidency of Strategy and Budget.

## Author contributions

**Conceptualization:** AV

**Design of experiments:** AV, ÖA

**Investigation:** ÖA, NA, ABK, SÜ

**Formal analysis:** AV, ÖA, SÜ

**Resources:** NA, ABK, TD, MÜ, IB, KS, MK, BY, AS

**Funding acquisition:** AV

**Visualization:** ÖA, SÜ

**Supervision:** AV

**Writing – original draft:** AV, ÖA, SÜ

**Writing – review & editing:** AV

## References

1. Baker D, Marta M, Pryce G, Giovannoni G, Schmierer K. Memory B Cells are Major Targets for Effective Immunotherapy in Relapsing Multiple Sclerosis. EBioMedicine. 2017 Feb;16:41–50.

2. Furman MJ, Meuth SG, Albrecht P, Dietrich M, Blum H, Mares J, et al. B cell targeted therapies in inflammatory autoimmune disease of the central nervous system. Front Immunol. 2023 Mar 9;14:1129906.

3. Hultin LE, Hausner MA, Hultin PM, Giorgi JV. Cd20 (pan-B cell) antigen is expressed at a low level on a subpopulation of human T lymphocytes. Cytometry. 1993 Jan;14(2):196–204.

4. Schuh E, Berer K, Mulazzani M, Feil K, Meinl I, Lahm H, et al. Features of Human CD3+CD20+ T Cells. J Immunol. 2016 Aug 15;197(4):1111–7.

5. von Essen MR, Ammitzbøll C, Hansen RH, Petersen ERS, McWilliam O, Marquart HV, et al. Proinflammatory CD20+ T cells in the pathogenesis of multiple sclerosis. Brain J Neurol. 2019 Jan 1;142(1):120–32.

6. Wilk E, Witte T, Marquardt N, Horvath T, Kalippke K, Scholz K, et al. Depletion of functionally active CD20+ T cells by rituximab treatment. Arthritis Rheum. 2009 Dec;60(12):3563–71.

7. Palanichamy A, Jahn S, Nickles D, Derstine M, Abounasr A, Hauser SL, et al. Rituximab efficiently depletes increased CD20-expressing T cells in multiple sclerosis patients. J Immunol Baltim Md 1950. 2014 Jul 15;193(2):580–6.

8. Ochs J, Nissimov N, Torke S, Freier M, Grondey K, Koch J, et al. Proinflammatory CD20 + T cells contribute to CNS-directed autoimmunity. Sci Transl Med. 2022 Mar 30;14(638):eabi4632.

9. Chen Q, Yuan S, Sun H, Peng L. CD3+CD20+ T cells and their roles in human diseases. Hum Immunol. 2019 Mar;80(3):191–4.

10. Mudd TW, Lu C, Klement JD, Liu K. MS4A1 expression and function in T cells in the colorectal cancer tumor microenvironment. Cell Immunol. 2021 Feb;360:104260.

11. Jonsson AH, Zhang F, Dunlap G, Gomez-Rivas E, Watts GFM, Faust HJ, et al. Granzyme K+ CD8 T cells form a core population in inflamed human tissue. Sci Transl Med. 2022 Jun 15;14(649):eabo0686.

12. Kaiserman D, Zhao P, Rowe CL, Leong A, Barlow N, Joeckel LT, et al. Granzyme K initiates IL-6 and IL-8 release from epithelial cells by activating protease-activated receptor 2. Chai KX, editor. PLOS ONE. 2022 Jul 26;17(7):e0270584.

13. Hsiao CC, Fransen NL, van den Bosch AMR, Brandwijk KIM, Huitinga I, Hamann J, et al. White matter lesions in multiple sclerosis are enriched for CD20dim CD8+ tissue-resident memory T cells. Eur J Immunol. 2021 Feb;51(2):483–6.

14. Kavaka V, Mutschler L, De La Rosa Del Val C, Eglseer K, Gómez Martínez AM, Flierl-Hecht A, et al. Twin study identifies early immunological and metabolic dysregulation of CD8 + T cells in multiple sclerosis. Sci Immunol [Internet]. 2024 Sep 27 [cited 2025 Jul 10];9(99). Available from: https://www.science.org/doi/10.1126/sciimmunol.adj8094

15. Wolf FA, Angerer P, Theis FJ. SCANPY: large-scale single-cell gene expression data analysis. Genome Biol. 2018 Feb 6;19(1):15.

16. Fang Z, Liu X, Peltz G. GSEApy: a comprehensive package for performing gene set enrichment analysis in Python. Lu Z, editor. Bioinformatics. 2023 Jan 1;39(1):btac757.

17. Xie Z, Bailey A, Kuleshov MV, Clarke DJB, Evangelista JE, Jenkins SL, et al. Gene Set Knowledge Discovery with Enrichr. Curr Protoc. 2021 Mar;1(3):e90.

18. Kuo WT, Kuo IY, Hsieh HC, Wu ST, Su WC, Wang YC. Rab37 mediates trafficking and membrane presentation of PD-1 to sustain T cell exhaustion in lung cancer. J Biomed Sci. 2024 Feb 7;31(1):20.

19. Vemulawada C, Renavikar PS, Crawford MP, Steward-Tharp S, Karandikar NJ. Disruption of IFN γ, GZM B, PRF 1, or LYST Results in Reduced Suppressive Function in Human CD8+ T Cells. J Immunol. 2024 Jun 1;212(11):1722–32.

20. Hamilton G, Colbert JD, Schuettelkopf AW, Watts C. Cystatin F is a cathepsin C-directed protease inhibitor regulated by proteolysis. EMBO J. 2008 Feb 6;27(3):499–508.

21. De Albuquerque R, Komsi E, Starskaia I, Ullah U, Lahesmaa R. The role of Interleukin-32 in autoimmunity. Scand J Immunol. 2021 Feb;93(2):e13012.

22. Lynch EA, Heijens CAW, Horst NF, Center DM, Cruikshank WW. Cutting Edge: IL-16/CD4 Preferentially Induces Th1 Cell Migration: Requirement of CCR5. J Immunol. 2003 Nov 15;171(10):4965–8.

23. Syed M, Dishman AF, Volkman BF, Walker TL. The multifaceted role of XCL1 in health and disease. Protein Sci. 2025 Feb;34(2):e70032.

24. Zeng Z, Lan T, Wei Y, Wei X. CCL5/CCR5 axis in human diseases and related treatments. Genes Dis. 2022 Jan;9(1):12–27.

25. Shiow LR, Rosen DB, Brdičková N, Xu Y, An J, Lanier LL, et al. CD69 acts downstream of interferon-α/β to inhibit S1P1 and lymphocyte egress from lymphoid organs. Nature. 2006 Mar;440(7083):540–4.

26. Lacotte S, Brun S, Muller S, Dumortier H. CXCR3, Inflammation, and Autoimmune Diseases. Ann N Y Acad Sci. 2009 Sep;1173(1):310–7.

27. Kondo T, Takiguchi M. Human memory CCR4+CD8+ T cell subset has the ability to produce multiple cytokines. Int Immunol. 2009 May 1;21(5):523–32.

28. Goedhart M, Gessel S, Van Der Voort R, Slot E, Lucas B, Gielen E, et al. CXCR4, but not CXCR3, drives CD8+ T-cell entry into and migration through the murine bone marrow. Eur J Immunol. 2019 Apr;49(4):576–89.

29. Mogilenko DA, Shpynov O, Andhey PS, Arthur L, Swain A, Esaulova E, et al. Comprehensive Profiling of an Aging Immune System Reveals Clonal GZMK+ CD8+ T Cells as Conserved Hallmark of Inflammaging. Immunity. 2021 Jan 12;54(1):99-115.e12.

30. Lan F, Li J, Miao W, Sun F, Duan S, Song Y, et al. GZMK-expressing CD8+ T cells promote recurrent airway inflammatory diseases. Nature. 2025 Feb 13;638(8050):490–8.

31. Guo CL, Wang CS, Wang ZC, Liu FF, Liu L, Yang Y, et al. Granzyme K+CD8+ T cells interact with fibroblasts to promote neutrophilic inflammation in nasal polyps. Nat Commun. 2024 Nov 29;15(1):10413.

32. Green DR, Droin N, Pinkoski M. Activation-induced cell death in T cells. Immunol Rev. 2003 Jun;193(1):70–81.

33. Kelly PN, Strasser A. The role of Bcl-2 and its pro-survival relatives in tumourigenesis and cancer therapy. Cell Death Differ. 2011 Sep;18(9):1414–24.

34. Kurtulus S, Tripathi P, Moreno-Fernandez ME, Sholl A, Katz JD, Grimes HL, et al. Bcl-2 allows effector and memory CD8+ T cells to tolerate higher expression of Bim. J Immunol Baltim Md 1950. 2011 May 15;186(10):5729–37.

35. McLane LM, Abdel-Hakeem MS, Wherry EJ. CD8 T Cell Exhaustion During Chronic Viral Infection and Cancer. Annu Rev Immunol. 2019 Apr 26;37(1):457–95.

36. Smith LK, Boukhaled GM, Condotta SA, Mazouz S, Guthmiller JJ, Vijay R, et al. Interleukin-10 Directly Inhibits CD8+ T Cell Function by Enhancing N-Glycan Branching to Decrease Antigen Sensitivity. Immunity. 2018 Feb 20;48(2):299-312.e5.

37. Chen Z, Ji Z, Ngiow SF, Manne S, Cai Z, Huang AC, et al. TCF-1-Centered Transcriptional Network Drives an Effector versus Exhausted CD8 T Cell-Fate Decision. Immunity. 2019 Nov;51(5):840-855.e5.

38. Schenkel JM, Herbst RH, Canner D, Li A, Hillman M, Shanahan SL, et al. Conventional type I dendritic cells maintain a reservoir of proliferative tumor-antigen specific TCF-1+ CD8+ T cells in tumor-draining lymph nodes. Immunity. 2021 Oct;54(10):2338-2353.e6.

39. Wu T, Ji Y, Moseman EA, Xu HC, Manglani M, Kirby M, et al. The TCF1-Bcl6 axis counteracts type I interferon to repress exhaustion and maintain T cell stemness. Sci Immunol. 2016 Dec 23;1(6):eaai8593.

40. Duquette D, Harmon C, Zaborowski A, Michelet X, O’Farrelly C, Winter D, et al. Human Granzyme K Is a Feature of Innate T Cells in Blood, Tissues, and Tumors, Responding to Cytokines Rather than TCR Stimulation. J Immunol. 2023 Aug 15;211(4):633–47.

41. Holley JE, Bremer E, Kendall AC, De Bruyn M, Helfrich W, Tarr JM, et al. CD20+inflammatory T-cells are present in blood and brain of multiple sclerosis patients and can be selectively targeted for apoptotic elimination. Mult Scler Relat Disord. 2014 Sep;3(5):650–8.

42. von Essen MR, Talbot J, Hansen RHH, Chow HH, Lundell H, Siebner HR, et al. Intrathecal CD8(+)CD20(+) T Cells in Primary Progressive Multiple Sclerosis. Neurol Neuroimmunol Neuroinflammation. 2023 Sep;10(5).

43. Hsiao CC, Engelenburg HJ, Rip J, Wierenga-Wolf AF, Van Puijfelik F, Van Luijn MM, et al. Acquisition of residency programs by T cells entering the human brain. Cell Rep. 2025 Jul;44(7):115960.

44. Quendt C, Ochs J, Häusser-Kinzel S, Häusler D, Weber MS. Proinflammatory CD20+ T Cells are Differentially Affected by Multiple Sclerosis Therapeutics. Ann Neurol. 2021 Nov;90(5):834–9.

45. Sabatino JJ, Wilson MR, Calabresi PA, Hauser SL, Schneck JP, Zamvil SS. Anti-CD20 therapy depletes activated myelin-specific CD8+ T cells in multiple sclerosis. Proc Natl Acad Sci U S A. 2019 Dec 17;116(51):25800–7.

46. Koetzier SC, van Langelaar J, Melief MJ, Wierenga-Wolf AF, Corsten CEA, Blok KM, et al. Distinct Effector Programs of Brain-Homing CD8+ T Cells in Multiple Sclerosis. Cells. 2022 May 13;11(10):1634.

47. Boldrini VO, Quintiliano RPS, Silva LS, Damasceno A, Santos LMB, Farias AS. Cytotoxic profile of CD3+CD20+ T cells in progressive multiple sclerosis. Mult Scler Relat Disord. 2021 Jul;52:103013.

