## Supplementary Figures for "CD8+CD20+ Cytotoxic T Lymphocytes Exhibit Augmented Degranulation and Pro-inflammatory Potential in Multiple Sclerosis"

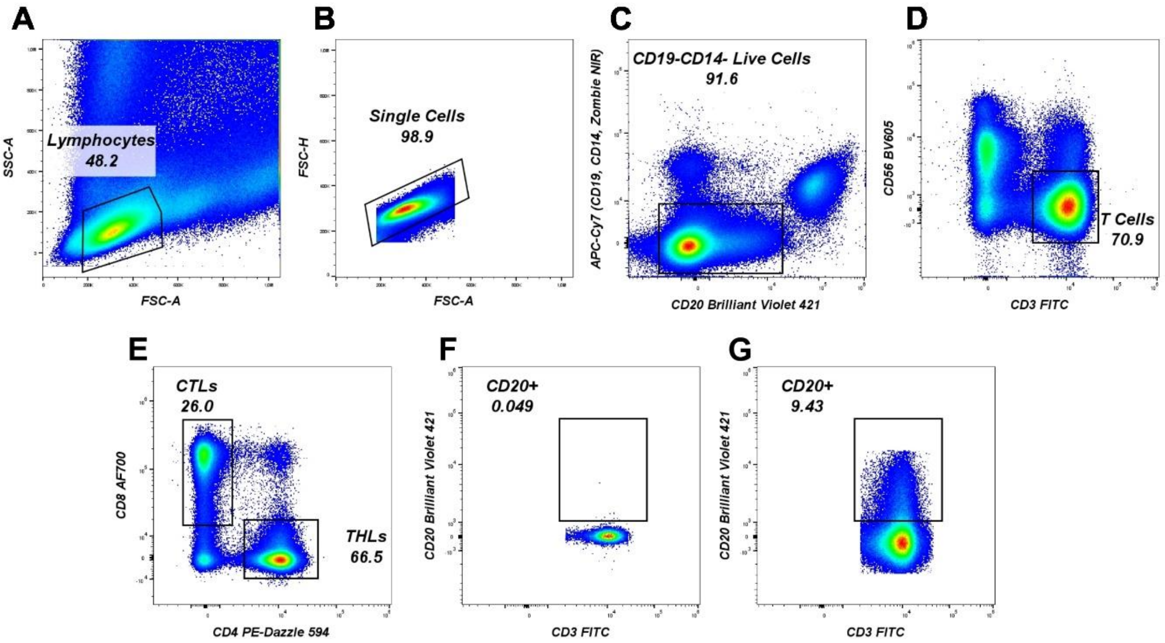


**Figure S1. F**igure illustrates the gating strategy for the analysis of CD20+ cytotoxic T cells. **A**) First, lymphocytes were selected from the FSC vs. SSC plot. **B**) Single cells were then gated from the lymphocyte gate to eliminate doublets. **C**) B cells, monocytes, and dead cells were excluded by plotting CD20 Brilliant Violet 421 against the dump channel APC-Cy7, which includes anti-CD19, anti-CD14, and Zombie NIR fixable viability dye. T Cels were selected as CD56-CD3+ cells **E**) CD8+ cytotoxic T cells were gated using the anti-CD4 PE-Dazzle 594 vs. anti-CD8 Alexa Fluor 700 plot. **F** and **G**) Finally, CD20 FMO was used to identify CD20+ cytotoxic T cells.

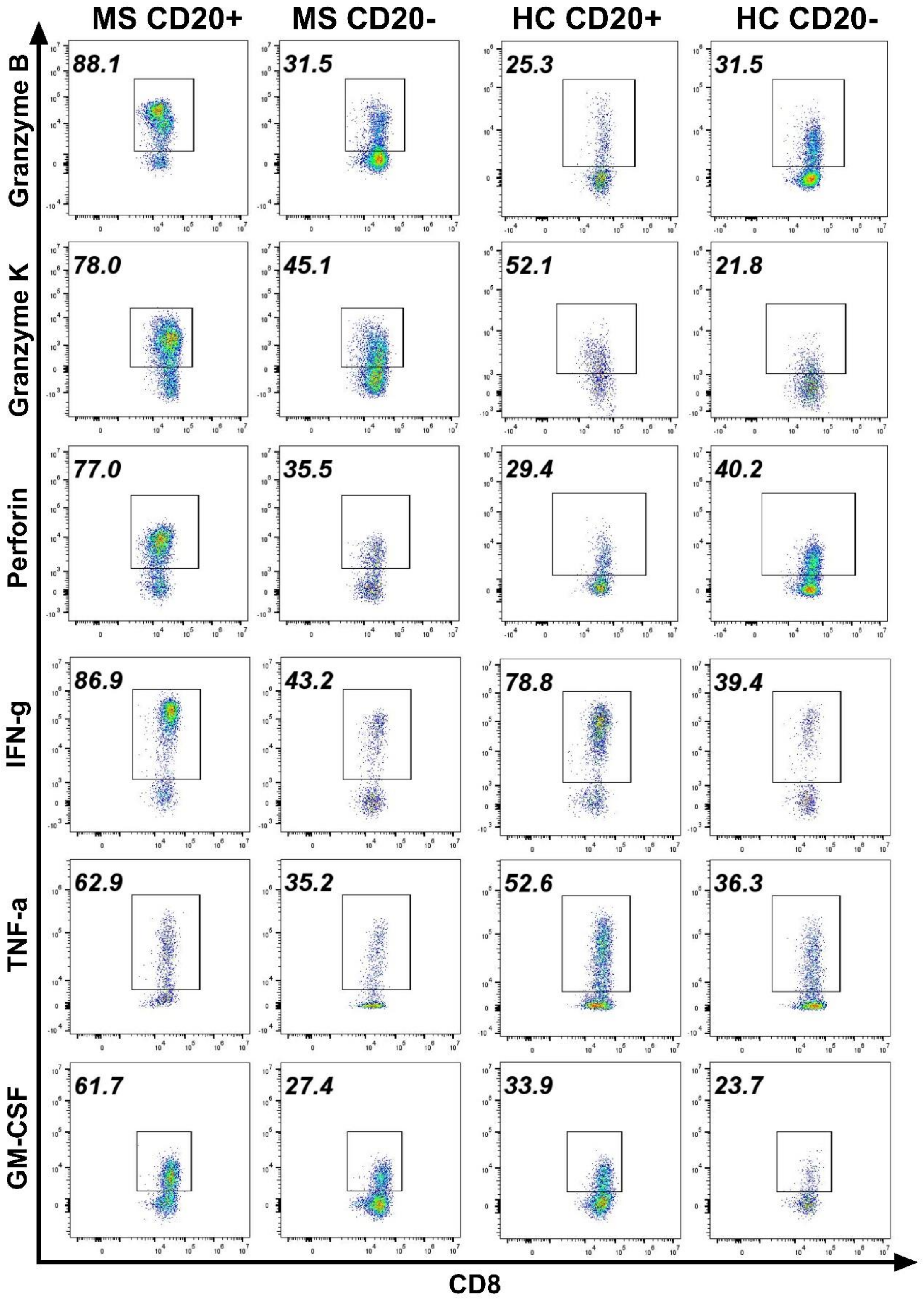


**Figure S2.** Figure shows the representative gating strategy for the functional analysis of CD20+ and CD20- CTLs in pwMS and healthy controls. Analysis was done on FlowJo Analysis Software v.10.10.0. Identical gates were created on both CD20+ and CD20- CTLs on a single sample and then the gates were applied to all samples ensuring that the same gating strategy was used for every sample.

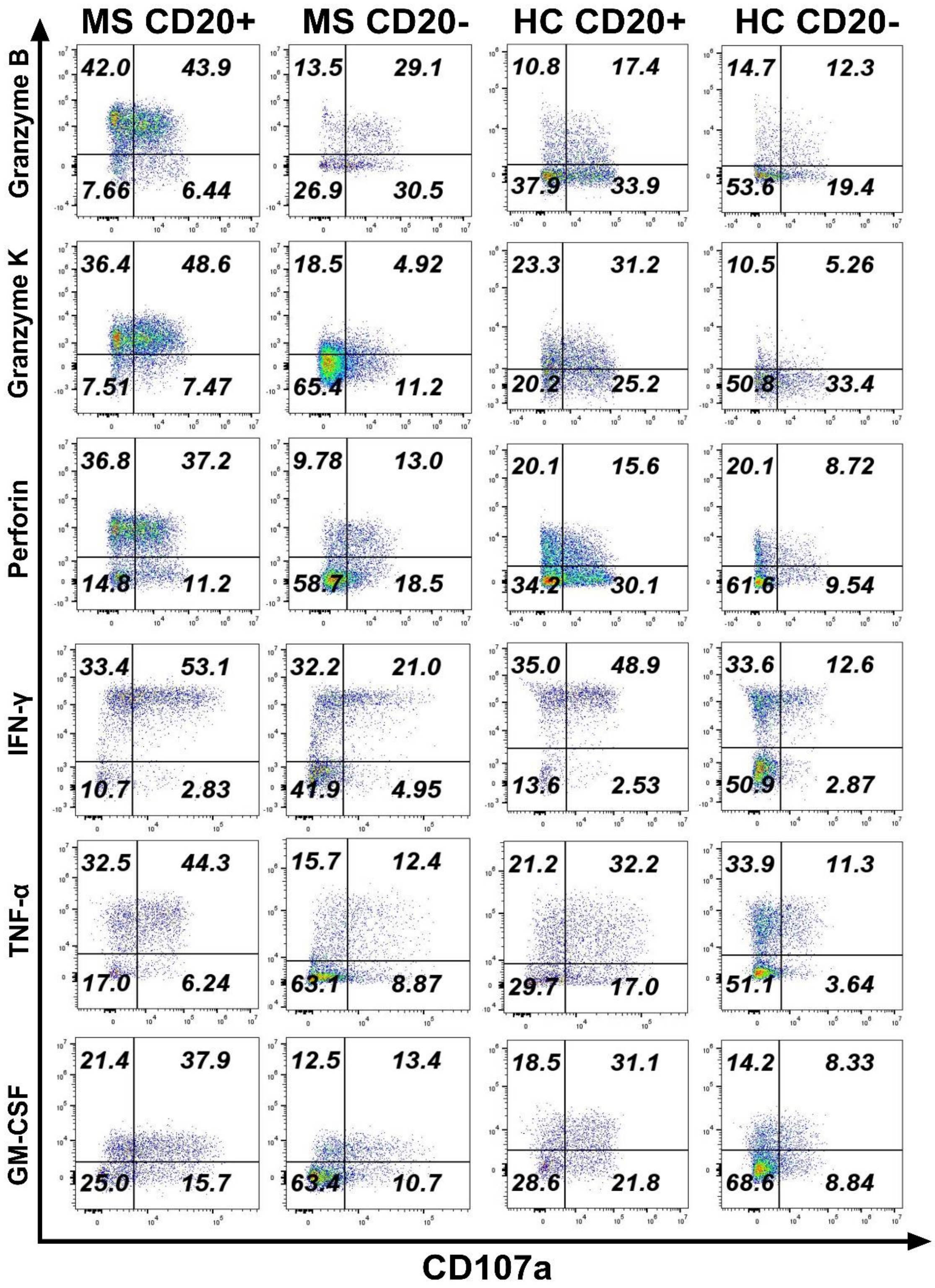


**Figure S3.** Figure shows the representative gating strategy for the analysis in CD107a+ population of CD20+ and CD20- CTLs in pwMS and healthy controls. Analysis was done on FlowJo Analysis Software v.10.10.0. Identical gates were created on both CD20+ and CD20- CD107a+ CTLs on a single sample and then the gates were applied to all samples ensuring that the same gating strategy was used for every sample.
